# Dynamic remodeling of fiber networks with stiff inclusions under compressive loading

**DOI:** 10.1101/2022.08.04.502849

**Authors:** Bobby Carroll, Minh-Tri Ho Thanh, Alison Patteson

**Affiliations:** Physics Department and BioInspired Institute, Syracuse University. Physics Building, Syracuse, NY 13244

**Keywords:** Compression stiffening of tissues, fiber networks with stiff inclusions, poroelasticity, network remodeling at the mesoscale, dynamic compression behavior

## Abstract

The ability of tissues to sustain and withstand mechanical stress is critical to tissue development and healthy tissue maintenance. The mechanical properties of tissues are typically considered to be dominated by the fibrous extracellular matrix (ECM) component of tissues. Fiber network mechanics can capture certain mechanical features of tissues, such as shear strain stiffening, but is insufficient in describing the compressive response of certain tissues and blood clots that are rich in extracellular matrix. To understand the mechanical response of tissues, we employ a contemporary mechanical model, a fibrous network of fibrin embedded with inert bead inclusions that preserve the volume-conserving constraints of cells in tissues. Combining bulk mechanical rheology and a custom imaging device, we show that the presence of inclusions alters the local dynamic remodeling of the networks undergoing uniaxial compressive strains and demonstrate non-affine correlated motion within a fiber-bead network, predicted to stretch fibers in the network and lead to the ability of the network to stiffen under compression, a key feature of real tissues. These findings have important implications for understanding how local structural properties of cells and ECM fibers impact the bulk mechanical response of real tissues.

## 1. Introduction

An early evolutionary feature of multicellular life is the production of an extracellular matrix that holds cell together and gives strength to multicellular structures. In mammals, cells develop into tissues. Tissues are comprised of both the volume-conserving cells and the cell-generated extracellular matrix, forming a dense composite cell-fiber network, permeated by interstitial fluid. There is much work demonstrating the properties of the extracellular matrix (ECM) determine the tissue mechanical properties^1^. With increased ECM deposition or crosslinking, cells increase tissue stiffness, and with increased ECM degradation, cells decrease tissue stiffness ^2^. Analyzing the properties of reconstituted fiber networks alone have revealed details of ECM stretching and stiffening under shear stress ^3^, which has been useful in describing shear strain stiffening of some whole tissues rich in collagen, such as the aorta^4^.

Recent work is showing however that ECM properties are not sufficient to predict tissue mechanical behavior, particularly under conditions of uniaxial compression. Most biopolymer networks soften under uniaxial compressive strain, whereas tissues stiffen ^5^. In traditional soft-condensed matter systems, a materials compression response is related to its shear response by the Poisson’s ratio v, which for isotropic linearly elastic materials follows the relation *G = E/2*(*1 + v*). But in tissues and the fibrous networks that form them, the Poisson’s ratio can be ill-defined and dependent on time scales and strain magnitudes in a manner not seen in most synthetic materials^6^.

Recent experiments have shown that reconstituted compression-softening fiber networks can be converted into compression-stiffening networks by the addition of volume-conserving beads ^5^, suggesting that tissue mechanics and rheology emerges from an interplay between the fibrous polymer network and the volume-conserving cells within them. To better understand the underlying mechanisms governing tissue mechanics, several models have been proposed to explain the emergence of this compression-stiffening in tissues and fiber-bead networks, largely focused on how volume-conserving beads might induce non-affine fiber stretching that stiffens the networks upon compression ^5, 7-8^, even at volume-inclusion densities well below jamming. Fiber stretching has been the suggested stiffening mechanism in other thermo-responsive fibrin-colloidal systems^9^. Mechanisms of compression-stiffening include biaxial fiber stretching around inclusions^5^, heterogenous relative motion of the inclusions ^8^, angle-constraining cross-linking and area conservation ^7^, and bead jamming upon compression^10^.

Traditionally, to build accurate computational or theoretical models of a material’s bulk to response to external forces, the material’s properties must be known at either at small scales, such as individual fibers (microscale); or at larger scales on the order of local clusters of interacting fibers (mesoscale). To validate or disprove these models, the fiber networks need to be experimentally probed at both the microscale and mesoscale. Confocal imaging of fibrin and collagen networks under compression have painted a more-complex picture of network response, with the emergence of a densified front near the compressing boundary^11-12^. Confocal and other advanced imaging techniques that can resolve single-fiber level configurations, however, largely do not have the time resolution to capture single fiber motions at the scale of whole-networks under compression in real-time. Thus, non-affine motions predicted to lead to fiber stretching and stiffening of the network ^13^ have not yet been captured in fiber-bead networks.

Here we present an experimental study of a model system to investigate dynamic uniaxial compression of soft fiber networks. We combine traditional rheology techniques that measure the bulk compression response of the material with a custom imaging device that allows us to examine the evolution of fiber and fiber-bead networks under compression at the meso-scale. In line with previous rheological investigations, the fiber-bead networks exhibit an apparent compression stiffening that is seen in native tissues, whereas the fiber network shows more network-like behavior under compression and does not stiffen. Analysis of the imaging compression videos reveals a complex mechanical response that proceeds through a series of remodeling processes, including formation of a densified front, lateral stretching in the network, and fluid fluxes through the network. From the analyses, a qualitative description of compression behavior for fiber-bead hydrogels is presented, potentially illuminating the mechanism of compression stiffening seen in native tissues.

Here, we experiment with two different types of networks: fiber and fiber-bead networks (Methods 2.1). Our fiber networks are 10 mg/mL fibrin gels, and our fiber-bead networks are prepared with the addition of volume-conserving, non-adherent dextran beads (50-150μm in diameter, shown in Figure 1a) at 20% volume fraction. As an experimental control, we include a very small fraction of dextran beads to the fiber networks (1%) to account for the difference in sample preparation due to the addition of the beads. Therefore, the fiber networks all have a small vol% of beads but behave as if they are simply fiber networks.

**Figure 1:**
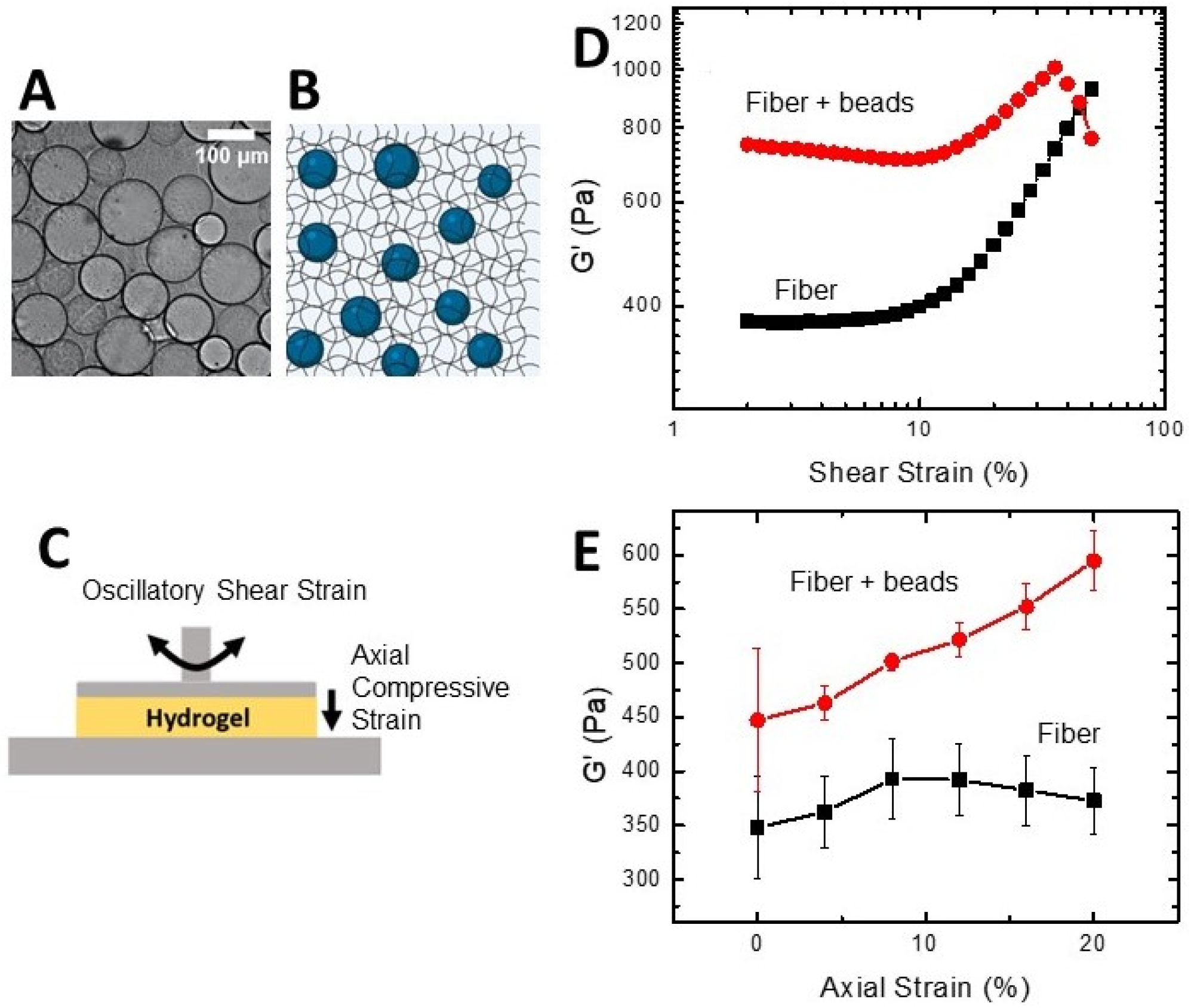
Fiber-bead networks exhibit compression-stiffening behavior. (a) A bright-field image of the fibrin network with embedded dextran beads (fiber-bead network). The dextran beads are 50-150 micron in diameter and clearly visible. (b) Schematic of fiber-bead hydrogel where a fibrin network is polymerized around the stiff dextran beads. The network is permeated by fluid that can move relative to the network. The mesh sizes are not to scale; the beads do not diffuse through the network. (c) Schematic of the rheology setup used to perform the compression tests. A continuous oscillatory shear strain is applied to measure the shear modulus *G* of the hydrogel at a starting gap height of 1mm. The axial compressive strain is controlled by successively lowering the gap height between the plates. (d) Representative plot of elastic shear modulus *G*□ measured for increasing shear strain amplitude for the fiber and fiber + beads cases. The fiber + beads case is 20vol% dextran beads. The beads modify the shear stiffening behavior expected from a fibrous polymer network (fiber). (e) The elastic storage modulus *G*□ taken after 5min of equilibration as a function of axial compressive strain for fiber and fiber-bead networks. The fiber-bead network stiffens upon increasing compressive strain. Each plot is averaged from three samples.

## 2. Methods

### 2.1 Fiber-Bead Network Formation

The model tissue hydrogels investigated are fibrin networks with stiff dextran beads embedded, as described previously ^5^. To prepare fibrin gels, fibrinogen (CalBioChem, EMD Millipore, Billerica, MA USA) was dissolved in T7 buffer (50mM Tris, 150mM NaCl, pH 7.4). Thrombin isolated from salmon plasma (seaRun Holdings, Freeport, ME, USA) is diluted in ddH_2_O at 1000 units per millimeter. Fibrinogen, thrombin, and CaCl_2_ were combined in 1X T7 buffer to yield a final solution of 10mg/ml fibrinogen, 30mM Ca^2+^ and 2 U ml thrombin.

Fibrin networks embedded with volume-conserving beads were prepared with cross-linked dextran beads (Sephadex G-25 medium, GE Health Sciences), as previously described ^5^. The beads are swollen with ddH20 to accomplish 92% swelling and are approximately 50-150 microns in diameter. Bead concentrations of 1vol% or 20vol% are used for the fiber hydrogels and fiber-bead hydrogels, respectively. To keep the concentration of fibrin, thrombin, and CaCl_2_ consistent, the extra volume % from dextran beads in the 20% case is accounted for by reducing the T7 buffer. The solution of fibrin, T7 buffer, CaCl_2_, and beads are pipette-mixed before thrombin is added to reduce the effect of sedimentation of the beads. Some of the T7 buffer (∼30mL) is added to the thrombin prior to mixing in the final hydrogel solution to prevent heterogeneous gel formation.

### 2.2 Rheology

All rheology tests were performed on a Malvern Panalytical Kinexus Ultra+ rheometer (Malvern Panalytical) equipped with a 20 mm parallel plate geometry. All samples were deposited on the bottom plate of the rheometer as solutions, immediately after initiation of polymerization. Then, the upper plate was lowered to a gap height of 1 mm. Gels were allowed to polymerize between the plates at 25°C, and the polymerization was tracked over time by measuring the sample shear modulus *G*□ and loss modulus *G*□□. The *G*□ and *G*□□ values were measured at 1 Hz and 2% shear strain during polymerization, which typically took 10 min to reach plateau values. Shear strain amplitude sweep tests were performed at 10Hz with shear strain amplitudes ranging from 0.2% to 50% shear stain. During compression tests, samples were subjected to small step-wise compressive strains, between which the samples were allowed to relax. The gap height between the parallel plates was sequentially lowered in steps of 4% compressive strain, reaching 20% strain, with 5-minute intervals for each step. All compressions tests were performed while simultaneously applying an oscillatory torque at 1 Hz and 2% shear strain. Additionally, creep recovery tests were performed on the fiber-bead networks. Creep tests were performed for 60s at 1Pa applied shear stress followed by 60s of recovery.

### 2.3 Compression Device

A custom-built compression device was made to image the mesoscale compression of the fiber-bead networks (Fig. 2a). Hydrogels are cast in the device between two glass slides for imaging, and compression of the sample is driven by a piezo motor (Newport NanoPZ ultra-high resolution motion system). The ramp speed of the compressing wall is 0.01mm/s. As viewed above, the gels are also confined above and below by two parallel glass walls, whereas the left and right sides of the gel constitute a gel-air interface and are free to move in response to applied deformation. The dimensions of the hydrogel in the imaging plane is 6.5 mm in width x 5-6 mm in length. The height of the sample in the z-direction is 1mm or approximately 10 bead diameters), which is thick enough to prevent buckling of the sample out of plane yet thin enough to allow transmitted light through the sample for microscopy imaging. The walls of the chamber are treated with Rain-x to reduce the sample sticking to the confining boundaries. The piezo motor pushes the compressing slide at a steady ramp of ∼0.2% strain/s up to 20% axial strain in two sequential videos: ∼0-20% (Video S1,S2) and ∼20-40%(Video S3,S4). After each ramp, the compression is stopped to allow enough time for full network relaxation (∼ 3 min).

**Figure 2:**
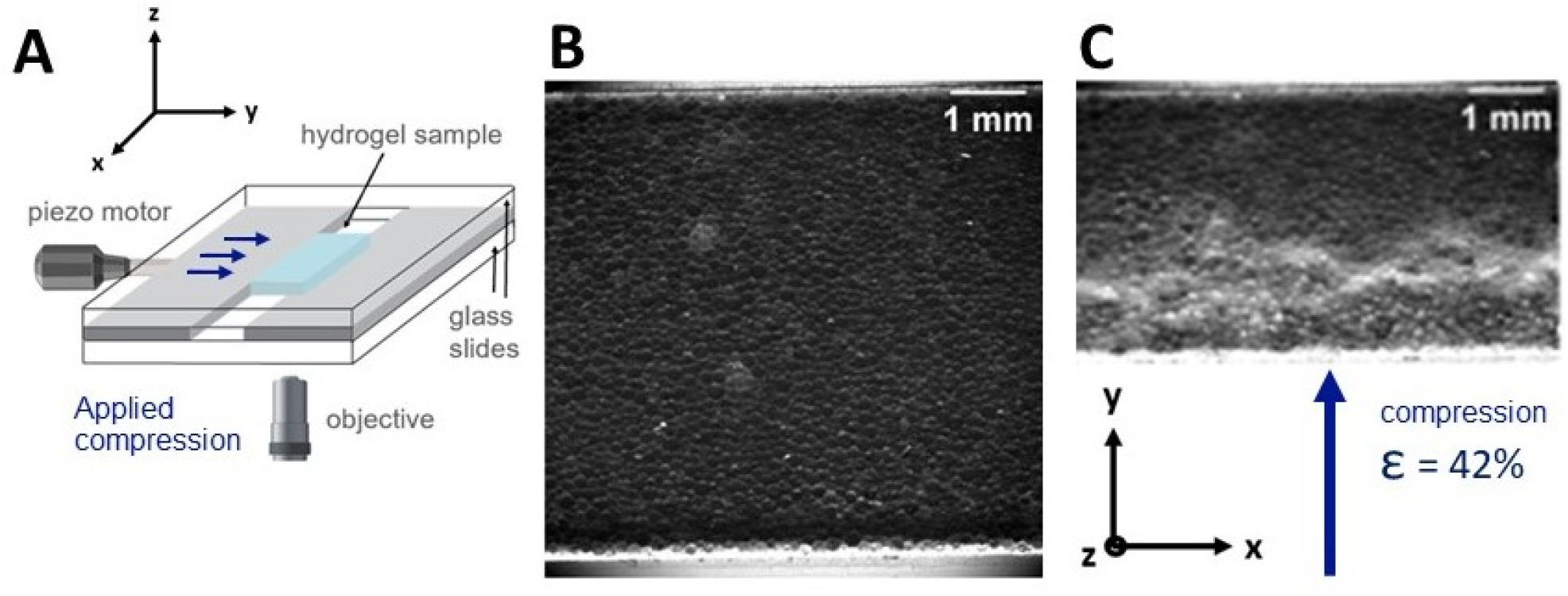
Experimental system. (a) Apparatus: custom compression device with imaging setup, with axes of direction. The fiber-bead hydrogel samples (blue) are polymerized in a transparent chamber between two glass slides (clear plates) and two walls (grey plates) for imaging. A piezo motor drives the moving wall (grey plate with blue arrows) in +y direction, compressing the sample against the stationary wall (grey plate without arrows). Liquid from the hydrogel is allowed to leave in the x-direction on either side of the sample. Representative phase contrast images of the (b) uncompressed and (c) compressed state of a fiber-bead network, where the direction of compression is in the +y-direction (axes have consistent orientation with (a)). The hydrogel sample deforms and rearranges, as it is compressed. A compression front emerges in the compressed sample, with a densification of beads near the driving wall.

### 2.4 Microscopy Imaging and Image Analysis

Fiber networks in the custom-built compression device are imaged with a Nikon Eclipse Ti (Nikon Instruments) inverted microscope, equipped with a 2x objective. Gels were imaged either in phase contrast or fluorescence; gels imaged with fluorescence are embedded with 5μm fluorescent polystyrene microspheres (Fluoro-Max, Part no. G0500) that function as tracers. Images were gathered at 1 frame per second with an Andor Technologies iXon em+ EMCCD camera (Andor Technologies). Images are captured either near the center of the sample or at the sample edge, where fluid flow out of the sample can be observed. All samples were imaged at room temperature. Fluorescence intensity profiles are taken by averaging the intensity values over the length of the x-direction for each pixel in y value, giving a plot of average intensity value vs length (mm in y-direction). To better compare intensity profiles for different frames, the intensity values are normalized to the area underneath the curve; an increase in intensity is assumed to correlate with an increase in fluorescent particle density. For the quantification of liquid leaving the gel, the area of the liquid is manually traced in Fiji for each frame, which multiplied by gap thickness gives volume. Tracing the liquid area can be accomplished from Videos S5,S6 by adjusting the brightness so the gel is oversaturated.

### 2.5 Particle Image Velocimetry

For imaging network deformations, 5 μm fluorescence microparticles (Fluoro-Max, Part no. G0500) were added to the fiber-bead networks. The fluorescence particle density is low enough (<0.01%) to not disrupt the network structure yet abundant enough to map deformations and relaxions of the network during compression. To obtain the network deformations, compression videos were analyzed with a custom particle image velocimetry (PIV) routine. This recursive routine employs a Fast Fourier Transform Cross Correlation (FFT-CC) algorithm provided by PIVlab ^14^ with a parallelization algorithm provided by ParaPIV^15^. The wrapper routine for verification uses a custom parallelization of Minimum Quadratic Difference algorithm provided by MPIV package^16^. Network velocity fields *v*(*r, t*) were extracted at approximately 100 μm intervals. Videos were obtained for both the applied compression and the network recovery, typically lasting 90 seconds for compression and 200 seconds for recovery. To quantify correlated deformations during the network recovery, we selected a central region of the network, at least 800 μm from the bottom-compressing and top-immobile walls, and computed the spatial velocity correlation function,

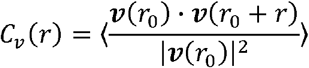

Here, the brackets denote an average over reference times *t*_0_ (in the range of 80 to 84 seconds after compression) and reference positions *r*_0_. Further, to isolate local correlated velocity within the bulk relaxation motion, we subtract out the average velocity in the window that we observe, before computing *C*_*v*_ (*r*). The high resolution deformation magnitude (color) heatmaps in Figure 4b,c were plotted by populating the 1000×1000 pixel grid with interpolated displacements from the nodal displacements (arrows) using MATLAB. The cumulative strain and shear maps in SI (Figure S6, S7) are calculated using formula given by Stamhuis et. al^17^: Shear 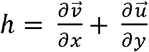 and Strain 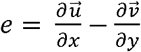. The 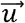 and 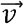 are horizontal and vertical components of the accumulated displacement at nodal position *x, y*. Accumulated deformation 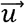 and 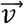 are calculated by summing up the pair-wise deformation fields of all time steps.^18^

**Figure 3:**
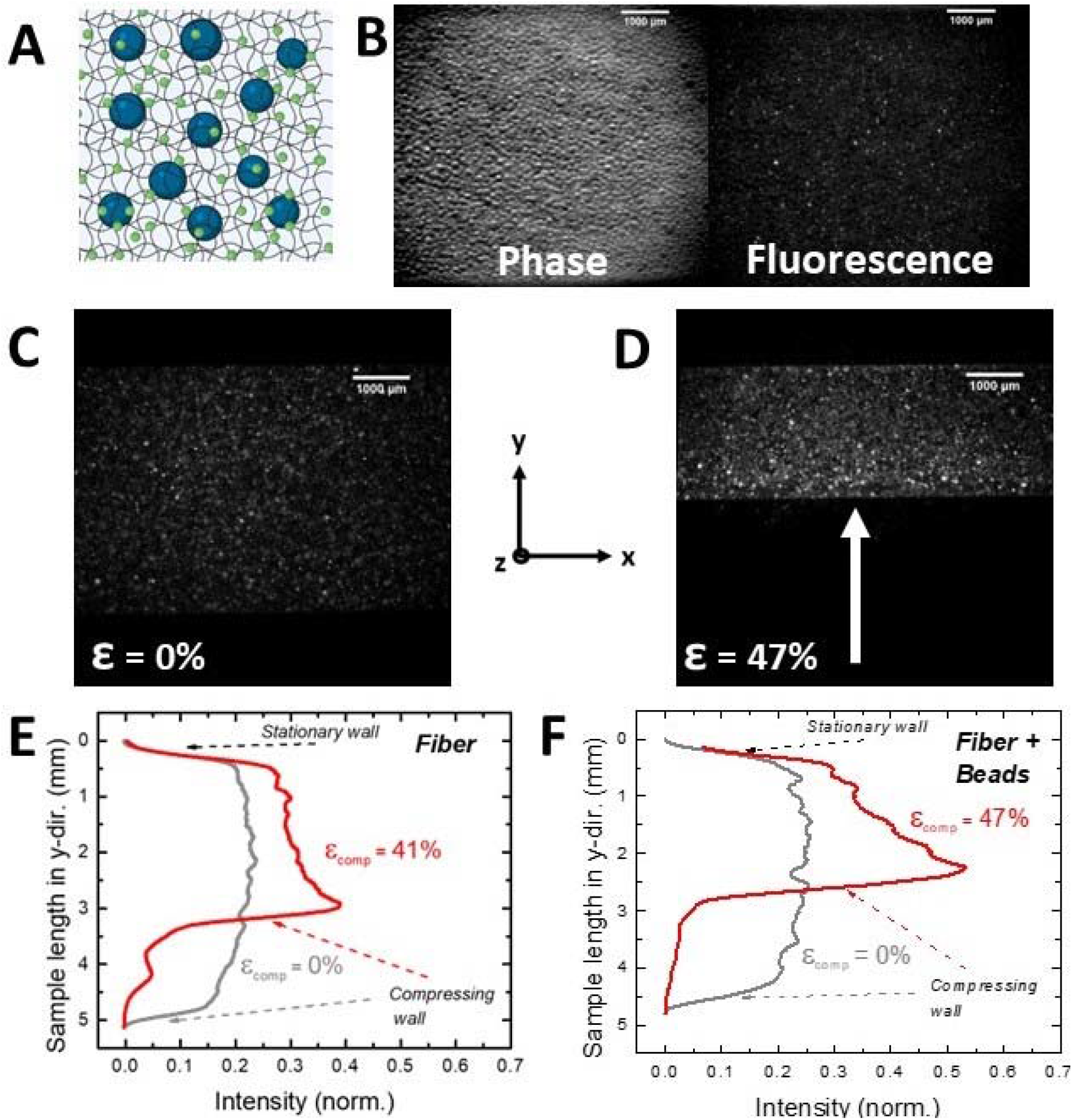
Local densification of fiber-bead networks upon compression. (a) To track the local deformation of the fiber-bead networks, small fluorescence tracers (green dots) were embedded in the network. The tracers are 5 μm in diameter, large enough to be trapped in the fibrin network. (b) Whereas the dextran beads are opaque and too dense to track over time, the fluorescence tracers are more dispersed, and their fluorescence signal is more amenable for tracking, as shown in sample phase contrast (left) and fluorescence images (right) of a fiber-bead sample. (c, d) Representative images of fluorescence tracers in a fiber-bead network upon compression. In the initial uncompressed (ε = 0%) state (c), the tracers are evenly dispersed throughout the sample. During compression (d), the fluorescent tracers accumulate near the compressing wall, indicating a densification of the fiber-bead network. (e, f) Quantification of network compaction. Plot of fluorescence intensity profiles across the length of the sample in the y-direction for fiber (e) and fiber-bead networks (f). The location of the stationary and compressing walls in the y-direction are indicated. The intensity profiles are shown for the initial uncompressed (grey) and compressed (red) state.

**Figure 4:**
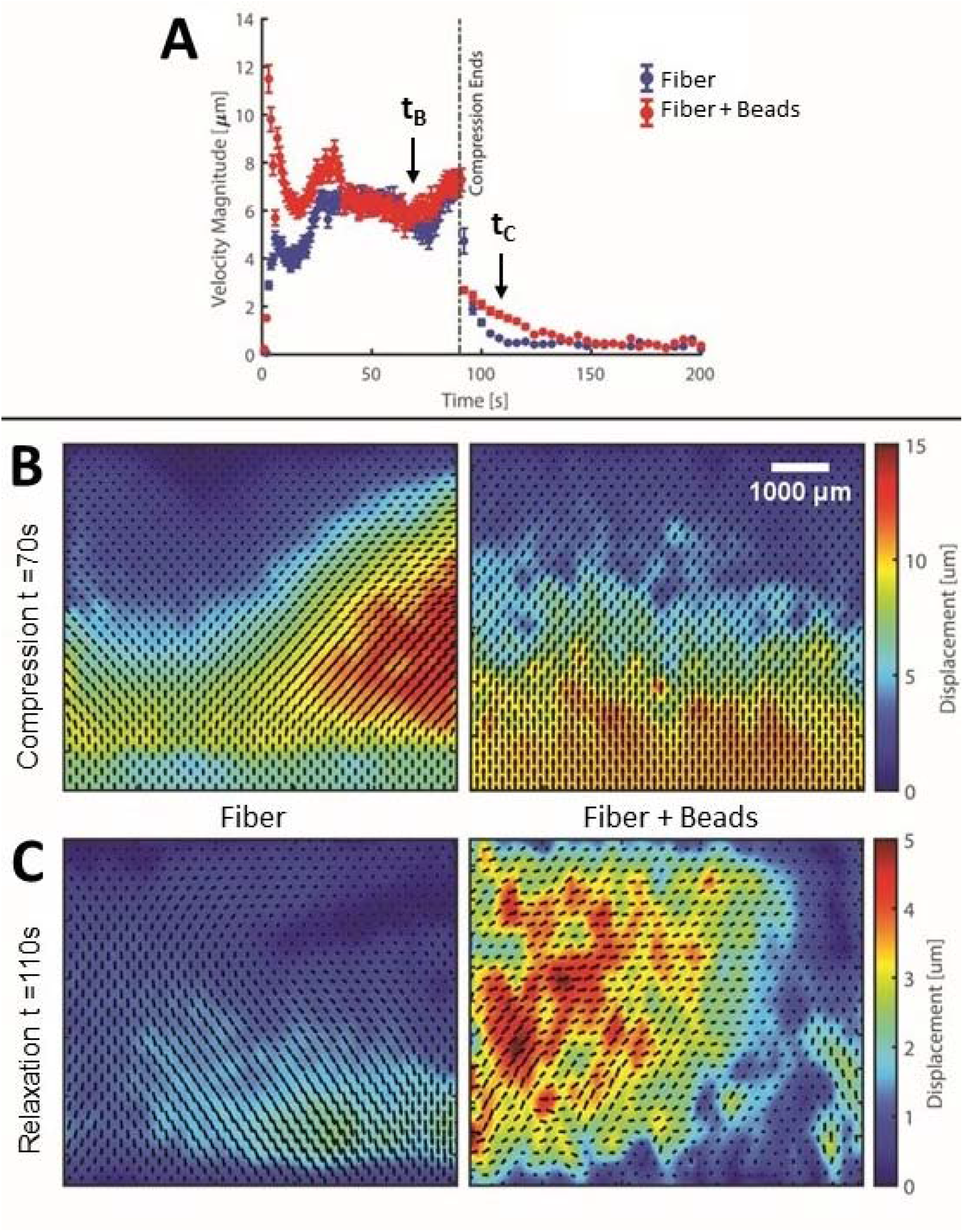
Network deformation fields. The local deformations of the network are measured over time via particle image velocimetry (PIV). (a) During compression, the samples exhibit global deformation with an average velocity magnitude of approximately 6 microns/second. When compression stops, the network relaxes, recovering from the applied load. The characteristic recovery time scale differs between the fiber and fiber-bead cases, with the fiber case recovering more quickly. The small arrows indicate the average velocity values at times *t*_*B*_ and *t*_*C*_ corresponding to the times the images in (b) and (c) are taken. The PIV velocity fields for the fiber case (left) and the fiber-bead case (right) are taken at (b) t = 70s (during compression) and (c) t = 110s (during relaxation). Each image is a heatmap of displacement magnitude from the previous frame (1s) with the corresponding velocity vector fields overlayed. Compression and relaxation have different displacement magnitude heatmap scales owing to their difference in velocity magnitude. Higher magnitude regions correspond to larger network deformation with time in that part of the network.

## 3. Results

Figure 1 shows the effect of volume-conserving beads (Fig. 1a, b) on the bulk mechanical response of fibrous networks from shear rheology. The hydrogel samples are loaded between two parallel rheometer plates (schematic in Fig 1c); fluid is free to leave the hydrogel radially from the gap, which is initially set to 1mm (full details in Methods 2.2). Subject to increasing oscillatory shear strain amplitude, the fiber-bead system has altered shear stiffening behavior in comparison to the fiber case (Fig. 1d). For the fiber-bead case, the G’ increases in magnitude at low strains and has less pronounced shear strain stiffening at higher strains; this nonlinear stiffening is more accurately described by the differential shear modulus *K* (Fig. S1). Very noticeably the fiber-bead network softens at high shear strain, indicating potential fiber breaking. Both fiber and fiber-bead networks have noticeable hysteresis with increasing then decreasing shear strain (Fig. S2). The volume-conserving beads also clearly affect the response to uniaxial compressive strain (Fig. 1e, Fig. S3). While an oscillatory torque of 2% shear strain and 1Hz frequency is applied, the hydrogels are subject to compressions of 4% steps up to 20% compressive strain. The data in Figure 1e shows that while the beads slightly increase the network’s shear storage modulus *G*□ in the initial uncompressed state, the beads have an even stronger effect on the network properties as the samples are compressed. Whereas the elastic shear modulus *G*□ of the fiber network is approximately constant upon compression, the fiber-bead network *stiffens* as it is compressed. The compression stiffening behavior observed here in fiber-bead networks also occurs in real tissues and collagen networks embedded with live cells^5, 19^, but is not seen in linearly elastic materials such as polyacrylamide gels or other fibrous networks, such as collagen^6^ or fibrin^20^ alone. The Poisson’s ratios calculated from the corresponding axial stress data (Fig. S3) do not correspond to linearly elastic networks. The compression behavior of the fiber network is similar to previous studies^5^ but there is less apparent softening. We finally note that the two systems exhibit compression hysteresis, as determined by measurements of G’ in the uncompressed state after the compression test was finished (Fig. S4); the fiber network had a reduction in stiffness (noticeable hysteresis) while the fiber-bead network did not.

To understand the bulk mechanical behavior of these networks, we examine the local rearrangement and deformations of the networks under uniaxial compressive loading using a custom-built device (Fig. 2). To image the local deformations, the networks are cast in a 1-mm thick layer between two glass slides (clear plates in Fig. 2a). The sample is thick enough to constitute multiple bead layers (∼10 bead diameters) yet thin enough to allow transmitted light for imaging with a microscope objective below the sample. The samples are subjected to a uniaxial compression strain via a piezo motor. Between the two clear glass slides, the sample is confined between two walls (grey plates in Fig. 2a). The top wall in the y-direction is stationary while the bottom wall (grey plate with blue arrows) is driven by the piezo motor, compressing the sample uniaxially at a slow, constant ramp speed *v* = 0.01 mm/s (Fig. 2a). Images taken from the device before and after compression are shown in Figure 2b, c. The imaging field of view for the uncompressed hydrogel is 6.6 mm x 6.2 mm (Fig. 2b), which for a fiber-bead network at 20% bead volume fraction allows imaging of approximately 4500 beads. Initially, the beads are evenly dispersed throughout the sample. As the driving wall compresses the sample in the +y direction, a densified front of beads emerges which travels along with compressing boundary. As seen in Figure 2c, the densified front is quite thick, approximately 1.9mm or ∼54% of the final sample height.

To track details of the fiber and fiber-bead network motion, we embed the networks with 5μm fluorescent tracers at small (<0.01%) concentrations (Fig. 3). The fluorescent tracers were much larger than the fibrin mesh size (∼100nm) ^21^ yet smaller than the dextran beads (50-150μm), such that motion of the fluorescence particles trace local mesoscale motions of the network. Figures 3 shows snapshots of the hydrogels used in the imaging experiment. (For full videos, see Videos S1-S4 of supplemental information). Each compression video contains a ramp compression of ∼90s followed by ∼200s of stationary imaging. Each network case has two of such videos, one from 0% - ∼20% compressive strain, and a second from ∼20-40% compressive strain (Full details in Methods 2.3). The videos of compression reveal three distinct features present in both the fiber and fiber-bead networks: (i) a global uniaxial compression accompanied by bulk lateral shear to the sides (x-direction), (ii) formation of a localized densified front near the compressing wall, and (iii) network recovery, where the network relaxes the applied stresses after the compression stops.

As shown in Figure 3, both the fiber and fiber-bead networks develop a densified front, as reflected by increased tracer density, but the thickness of the front depends on the presence of the volume-conserving dextran beads. The densified phase can be identified by increased fluorescence intensity in the images, reflecting regions of increased network density. To quantify the densified fronts, we computed the average image intensity profiles *ø(y,t)*, where y is the distance along the direction of compression between the compressing and stationary walls (Fig. 3e, f). As shown in Figure 3, the presence of the dextran beads noticeably alters the intensity profile of the fiber network. The front appears to extend further away from the compressing wall as compared to the fiber network case. To quantify this effect, we identify the location of the densified font boundary as the distance from the peak in intensity to where the intensity was equal to the average between the peak intensity value and the intensity near the stationary wall. This calculation shows the densified front went ∼0.4 mm for fiber networks to ∼0.7 mm in the fiber-bead network. These compressed fronts are consistent with conditions when the fibrin networks contain a fraction of fluorescently labeled fibrin, instead of fluorescence micro-particles (Fig. S5).

Similar densified fronts in fiber networks have been previously reported ^11-12, 22^. It has been argued that the densified front is induced by local fiber buckling under compression and alignment of fibers perpendicular to the stress ^11-12, 22^. Densification has also been associated with the unbinding of fiber crosslinks driven by cell contractile forces^23^. Granular systems have also been reported to have densified fronts, where uniaxial compression leads to a dynamic jamming front ^24-25^. In our experiments, the fiber networks are combined with stiff inclusions (beads), and the volume fraction of the beads (20%) is much lower than the volume fraction for jamming in 3D (64%^26^). Here, we find the inclusion of beads noticeably increases the extent of the densified front. One possible explanation is that the inclusion of stiff beads in the network at least partially interrupts the alleviation of stress, which occurs via fiber buckling and alignment in a narrowly concentrated zone, such that the densified region must extend further away from the wall. Within the densified front, we expect the fibers are buckled, stretched, or possibly broken, with the latter two more likely in the fiber-bead network.

Next, we examined the network deformation fields upon uniaxial compression and subsequent network recovery via particle image velocimetry (PIV, Methods 2.5). Figure 4 shows sample velocity fields in both the fiber and fiber-bead networks (Fig. 4b,c). The heatmaps demonstrate the change in displacement magnitude from the previous frame (1s), with the corresponding velocity vector fields overlayed. As shown in Fig. 4, the speed of the tracer particles near the moving wall are close to the speed of compressing wall *v*, which is expected for uniaxial compressive loading. We also observe significant lateral stretching in the samples during compression. This lateral stretching is the shearing motion of the sample in both x directions which varies across the plane of compression. This lateral stretching can be seen in Fig. 4b, where the imaging window is positioned near the center of the sample. The clearest demonstration is seen dynamically in Videos S1, S2 for the fiber and fiber-bead networks, respectively. The network deformation can also be presented with the cumulative strain and shear maps (Fig. S6, S7), which shows directional deformation throughout the network accumulated over the course of the compression and relaxation, respectively.

When compression stops, we observe a time-dependent relaxation of the network. The network creeps back, relaxing the stresses applied at the boundary throughout the network (SI Videos 1 and 2 for fiber and fiber-bead networks, respectively). Figure 4a shows the average velocities of the particles during the compression and relaxation process for the fiber and fiber-bead networks. During compression, the average velocity is relatively constant, driven by the displacement of the compressing boundary. Once the compression ramp stops, the networks creep back with an average velocity that decays toward zero over a characteristic time. Interestingly, the velocity magnitude decay is significantly longer in the fiber-bead network as compared to the fiber network. To obtain a characteristic network relaxation time *τ*_*PIV*_, we fit the PIV-average velocity decay curves to *v* = *v*_0_ exp (− (*t*− *t*_0_)/ *τ*), where *v*_0_ is the average velocity during compression and t_0_ is the time when the compression stops. The presence of the dextran beads increases the relaxation time *τ*_PIV_ more than 3-fold, with 12s for fiber network compared to 41s for the fiber-bead network (Table 1).

**Table 1:** Summary of timescales relevant to the relaxation of the fiber networks with stiff inclusions. Each timescale has a notation in column 2. Normal force *N* and elastic shear modulus *G*□ decay times are determined from the rheology compression tests. The time scales are shown are for the first compressive step at 4% axial strain, but similar values were obtained for the subsequent compressive steps. The other rheometer timescale is from the recovery following a constantly applied strain (creep recovery test). The PIV Velocity timescale is extracted from the decay in average velocity with time from particle imaging velocimetry analysis. Finally, the fluid flow timescale comes from the change in volume with time of the liquid flowing out of the edge of the hydrogel. All error bars are standard error of the mean, with the number of samples indicated in the first column.

| Timescale (Method) | Timescale notation | t (s) for fiber networks | t (s) for fiber-bead networks |
| --- | --- | --- | --- |
| Normal force decay (Rheometer) N = 3 | $\tau_N$ | 48±1s | 34±5s |
| $G'$ decay N =3 | $\tau_{G'}$ | 71±2s | 59±5s |
| Creep and recovery (Rheometer) N = 3 | $\tau_{CR}$ | 16±1s | 19±5s |
| PIV velocity decay N = 2 | $\tau_{PIV}$ | 12±6s | 41±13s |
| Fluid flow out of hydrogel edge N = 2 | $\tau_{FF}$ | 118±4s | 58±9s |

The two-dimensional (2D) velocity maps shown in Figure 4b and 4c also reveal details of the relaxation activity. In the absence of volume-conserving beads, the sample deformations smoothly vary throughout the sample, with some deviations near the boundaries of the sample. Interestingly, in samples with volume-conserving beads, we observe local collective motions throughout the sample. To quantify these collective motions, we compute the 2D velocity autocorrelation function *C*_*v*_(*r*) over a short time window (4s), where we have subtracted out the mean relaxation velocity in the observed portion of the hydrogel (Fig. 5a, b; Methods 2.5). While the *C*_*v*_(*r*) data is relatively flat for the fiber network, a clear peak arises in the fiber-bead network. At small r (or within a region of collective motion), the local velocities are correlated and *C*_*v*_(*r*), is relatively high. As r increases and extends past the length scale of correlated motion, the magnitude decays. As shown in the radial average of the velocity autocorrelation data (Fig. 5c), the correlated motion in the fiber-bead network has a correlation length of approximately 0.7 mm, which is ∼7 dextran bead diameters. Taken together, Figures 4 and 5 suggest the presence of stiff volume-conserving beads induces heterogeneous rearrangements within the sample, impacting the relaxation modes of the network due to uniaxial compression.

**Figure 5:**
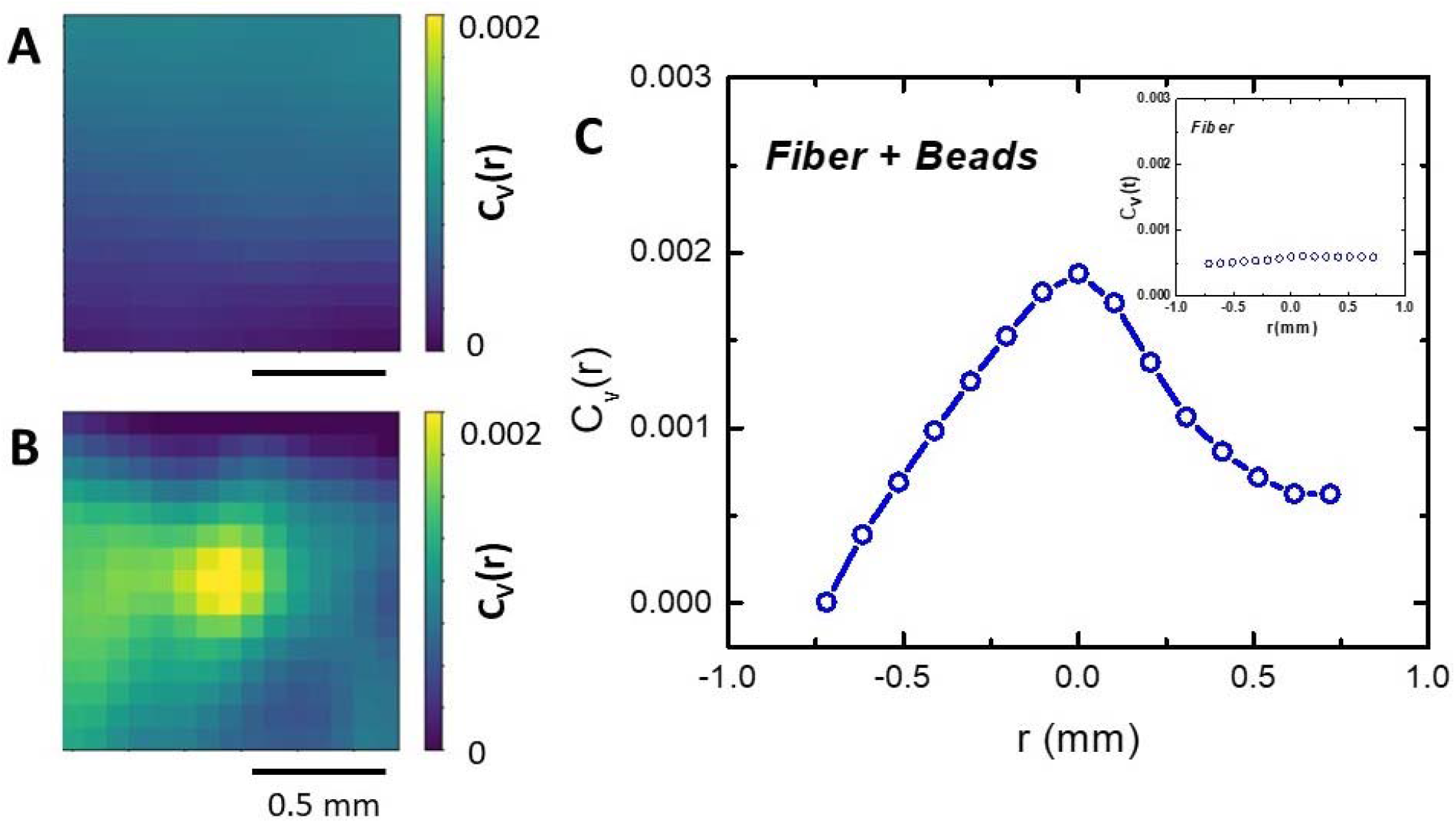
Localized rearrangements of the network under compressive deformations. (a) The magnitude of the velocity autocorrelation function for (a) fiber and (b) fiber-bead networks in 2D over a short time window (4s). The correlation data is from a central region of the hydrogel, after the mean relaxation velocity has been subtracted out. The yellow peak in the center at r=0 indicates strong correlated motion in the fiber-bead sample. (c) Velocity correlation coefficient as a function of r for the fiber + beads case. The correlated motion in the fiber-bead networks has a correlation length of ∼0.7mm. (Inset) The velocity correlation coefficient for the fiber network, which demonstrates no correlated motion.

Taken together, our data shows that volume conserving inclusions impact the mesoscale structure and dynamics of fiber networks, and these changes correlate with the onset of compression stiffening behavior seen in macroscopic rheological characterization. In particular, we find that the presence of the dextran beads increases the extent of the densified front, thereby increasing its reach into the sample (Fig. 3e, f). Further, the beads induce long-scale correlated rearrangements upon network relaxation (Fig. 5a, b).

Based on our results, we posited that the presence of volume-conserving beads alters the time-dependent relaxation of the network at the global scale (hydrogel scale), which could be measured via macroscopic rheology tests. To better understand the relaxation dynamics of the networks, we examined the response of the networks to two modalities of applied stress: uniaxial compression and creep recovery experiments, seen in Figure 6.

**Figure 6:**
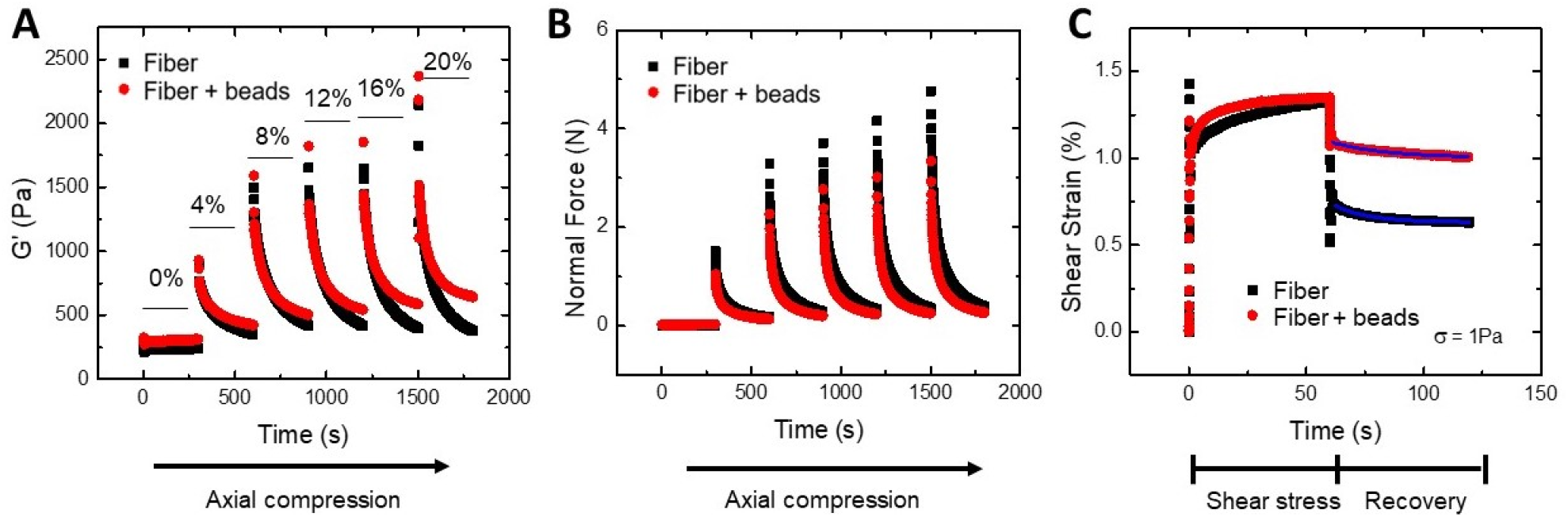
Network response to applied stress from bulk shear rheology. (a) The transient storage modulus measured during the compression tests from bulk shear rheology. The compression test measures the storage modulus for five minutes per step, where each curve represents the response after a 4% compressive strain is applied, up to 20% compression. (b) The transient normal force response is shown for each 4% compression step. This data is from the same compression test of the corresponding storage modulus data in (a). The decay in normal force appears to happen faster for the fiber-bead network than the fiber network alone. (c) Creep and recovery experiments for the fiber and fiber-bead network cases. The fiber-bead network has less strain recovery after the applied shear stress of 1Pa is removed. The recovery is fit to a single exponential decay (blue lines); the decay time is slightly longer for the fiber-bead case than the fiber alone.

The evolution of the normal force *N* (Fig. 6a) and elastic shear modulus *G*□ (Fig. 6b) shows details of the sample relaxations under a series of uniaxial compressive steps of 4% axial strain. Upon each compressive step, both the normal force and elastic shear modulus shoot up and subsequently decay to a steady state value with a characteristic time scale. Upon compression, the steady state values of *G*□ differ between the two network types. The relaxation of *G*□ for the fiber network alone consistently decays toward its value in the uncompressed state, whereas the fiber-bead network becomes stiffer with each compressive step, consistent with the compression stiffening behavior shown in Figure 1. Unlike the elastic shear modulus *G*□, the normal force steady state values were much more similar between the two gel types.

To characterize the relaxation dynamics, we fit both the normal force curves to a single exponential decay function of the form *f(t)* = *f*_0_exp (−*t / τ*) to obtain a characteristic time scale *τ*_*N*_ (Table 1). Surprisingly, we found that the dynamic relaxation of networks was faster in fiber-bead network compared to fiber network under uniaxial compression. We also measured the relaxation time scale based on the *G*□ curves, *τ*_G′_ (Table 1). These timescale values were consistently higher than those of the normal force decay; nonetheless, they followed the same trend of the fiber-bead network decaying faster than the fiber network. The relaxation of the networks contains components both from viscoelasticity and poroelasticity, where the poroelastic contribution is contact area dependent and the viscoelastic contribution is not. The axial stress relaxation changes with contact area (Fig. S8), indicating that poroelasticity is an important contribution to relaxation for both fiber and fiber-bead networks.

Since uniaxial compression induces large lateral stretching in our samples (Fig. 4; Video S1-S4), we also examined the network response to applied shear stress via creep recovery tests. As shown in Figure 6c, the fiber-bead samples recovers significantly less than the fiber networks alone after an applied shear stress of 1Pa. Plasticity is a feature of fibrin networks that is understood to be caused by network remodeling^27^, suggesting the beads facilitate more significant fiber remodeling during shear. To characterize the network relaxation dynamics from an applied shear stress, we fit the recovery portion of the curve to a single exponential decay to obtain a characteristic recovery time scale *τ*_*CR*_. The recovery time of the fiber-bead network (19 sec) was slightly larger in response to shear stress than the fiber network (16 sec), showing the fiber networks relaxed more quickly in response to applied shear strain (Table 1). Intriguingly, this result was opposite the relaxation dynamics in response to uniaxial compression, where the fiber-bead networks relaxed more quickly (Fig. 6a, b; Table 1).

## 4. Discussion

What might cause these differing effects of stiff inclusions on network dynamics, and how do network relaxation modes impact the compressive stiffening behavior of fiber networks? To interpret our results, we suggest a minimal model in which uniaxial compression sets up two different but connected relaxation process in fiber networks, which act as poro-elastic materials^28-30^. In this picture, the network behaves as a poroelastic material permeated by fluid that can move relative to this network. First, given the incompressibility of water, it would stand to reason the dominant cause of the normal force rise and decay upon uniaxial compression would be fluid pressure. As the network is compressed, the pressure rises sharply; this induces pressure gradients that drive the fluid through the network. As more of the fluid leaves the network, the network bears more of the applied load, relaxing the normal force on the bounding plate over time^29^. Second, uniaxial compression drives large lateral stretching of the network (Fig. 4)^8^. This imposes large shear stresses throughout the network, which we speculate stretch individual fibers in the direction perpendicular to compression. The relaxation of the network thus depends in part on the relaxation of the stretched fibers and on fiber-fiber interactions, described by a creep recovery process. We note that in a poroelastic material, transient stresses can set up fluid flows and network remodeling, such that these two processes are coupled and not entirely independent^31^. Previous studies have described the complex interplay between fiber remodeling and outgoing solvent flow on the nonlinear behavior of biopolymer gels20, 32-33.

Motivated by our findings, we propose that the main features of uniaxial compression in these fiber networks are normal force relaxations dominated by poro-elastic fluid flow and fiber stretching from lateral shear. The shorter normal force relaxation for the fiber-bead network case implies the fluid is leaving the sample faster than in the fiber network alone case. As we and others have shown, the presence of the dextran beads stiffen the network^5^ under compression owing to bead volume preservation which subsequently causes fiber stretching. Further, for poroelastic materials, we expect a stiffer network transmits greater stress through the material, driving faster fluid flows as compared to a soft network. Faster fluid flow has been associated with stiffer networks previously^34-35^. In addition, the network motion captured in the imaging device demonstrates that hydrogels undergo internal lateral shearing in response to compression and a subsequent relaxation. Our creep recovery test shows that the fiber-bead network recovers less and at a slower time scale *τ*_*CR*_ compared to the fiber network alone (Fig. 6c), which is consistent with PIV-observed relaxations (Fig.4). These results indicate that the presence of dextran beads hinders network recovery to applied shear stresses and the network holds larger internal stresses, which cannot be relaxed. These internal stresses stiffen the network^8^, consistent with the observed compression-stiffening behavior in the bulk rheometer (Fig. 1). While the bulk normal force relaxation is dominated by the fluid pressure, the effect of the dextran beads on network can be detected by its effect on the elastic shear modulus *G*□.

This argument, however, is contingent on the expectation that the networks expel fluid out of the network at timescales relevant to normal force decay. To gain further insight and verify that this is the case, we directly visualized the edge of our hydrogel samples during compressive loading in the imaging device and visualized flow of fluid leaving the network (Fig. 7, Video S5, S6). After compression starts, the network buckles slightly before the edge of the network breaks and fluid begins escaping the network (Fig. 7a, Video S5, S6). The once compression stops, the network relaxes back towards the center (Fig. 7biii) but the liquid continues leaving the edge, albeit at a slower pace. Figure 7c shows the volume (in μl) of the fluid leaving the edge of sample over time for both the fiber and fiber-bead networks (see Methods 2.4 for volume measurement method). We find indeed that rate of fluid flow escaping the network is higher in the fiber-bead network than the fiber case. This can be quantified with an exponential association function of the form *f(t*) = *f*_0_(1 − exp(−*t*/*τ*)); the timescale *τ*_*FF*_ for the fiber-bead network is significantly shorter (58 s) compared to the fiber network alone (118 s). We note that these timescales are notably larger than the timescales for normal force decay TN in the rheometer (Fig. 6b, Table 1), but this is expected because fluid must travel farther to escape the network in the compression device than in the rheometer given its boundary constraints.

**Figure 7:**
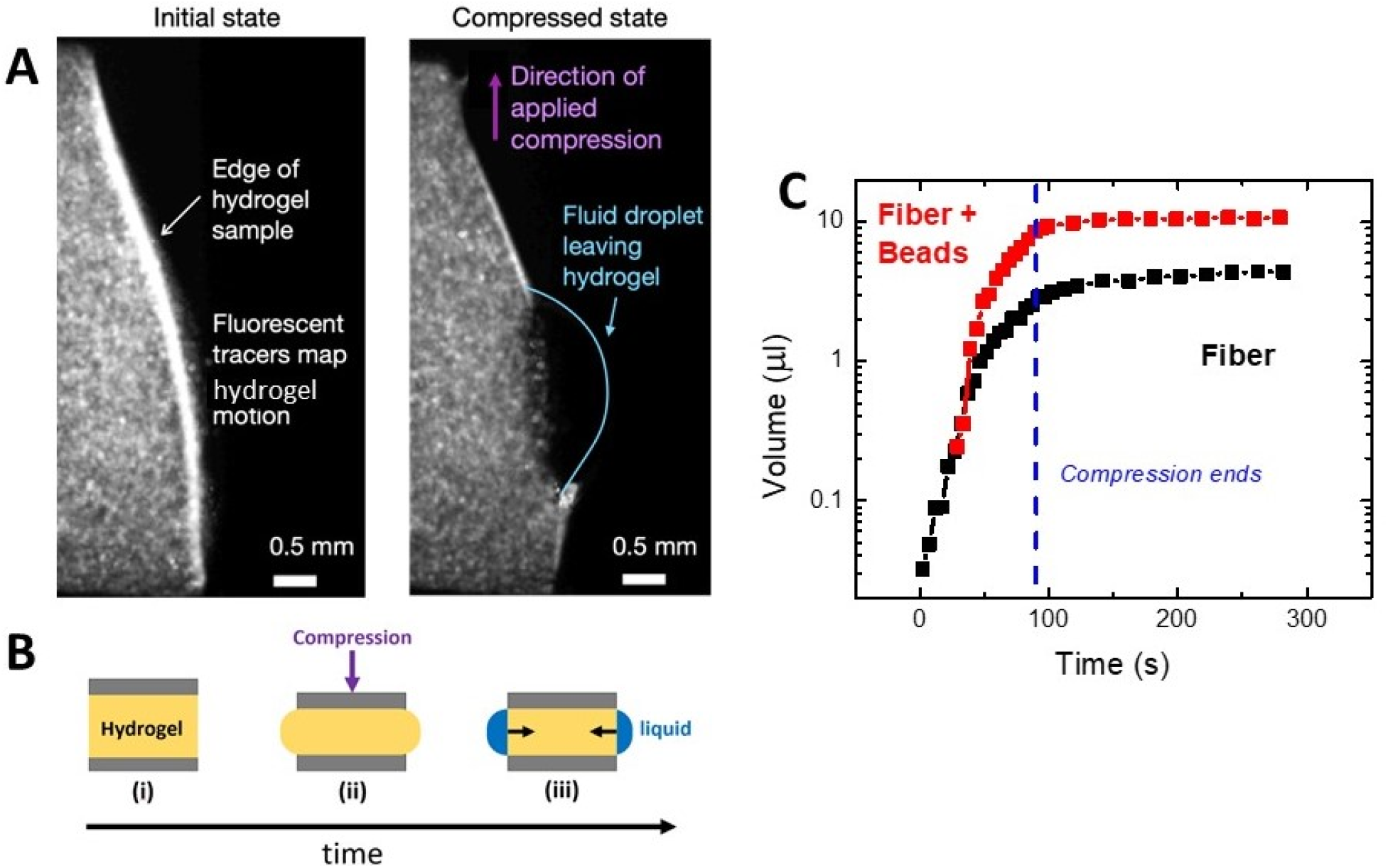
Poroelastic fluid flows upon compression. (a) Representative images of the network edge during compression. The network contains a percentage of fibrin which is fluorescently labelled. Immediately following the compression, the network edge bulges in response to the compressing force. After a few seconds of compression, the edge breaks and the fluid flows out the network (b) Schematic of sample compression and resulting fluid flow. The initially uncompressed hydrogel (i) has a compressive strain applied for a given amount of time. The edge bulges initially (ii) during compression before breaking, allowing liquid to flow. Once compression stops (iii), the network relaxes back towards the center (black arrows) while liquid continues to flow. (c) The volume of liquid leaving the edge of the sample as a function of time. Fluid is expelled more quickly from the fiber-bead network than the fiber network alone. More total fluid is expelled the fiber-bead case as well. Fluid continues to flow out the edge after compression stops at a slower pace.

Further, the available surface area for liquid to escape is larger for the rheometer than the imaging device, ∼63mm^2^ vs ∼10mm^2^, respectively. The compression procedures are different as well; a steady ramp for the imaging device while the rheometer has successive compression jumps with equilibration time in between. We expect that these timescales would be the same if the two experimental systems had equivalent geometry and compression procedures. To reiterate, the effect of dextran beads in the network increases the rate of fluid flow out of the network, which is consistent with the more rapid normal force relaxation in the rheometer (Fig. 6b, Table 1).

Further studies on this fiber-bead network should investigate the role of compression rate on deformation behavior. The steady ramp studied here in the visualization experiments is quasi-static, and the interplay between fiber network relaxation and fluid flow would be very different for a faster compression rate. Additionally, exploring how the deformation changes with thicker samples in the z-direction would give a more 3D description of compression behavior and would be more relevant to real tissue systems. Finally, tuning concentration of fibrin and dextran beads would give insights to how tissues of differing biopolymer and cell concentrations might behave in real tissues.

We emphasize that the fiber-bead networks investigated here are only a model system that captures some features of native tissues. The role that active cell forces as well as extracellular matrix proteins like proteoglycans are not included in this description of compression stiffening. Cells in native tissues are more deformable and smaller than the dextran beads used in this system, so while they will retain compression stiffening the fiber network deformation may be qualitatively different. Processes like cell migration also ensure there is an active component of tissue mechanical properties not captured in the equilibrium system here. A further study may investigate the behavior of cells under compression in these fiber network structures. Cell orientation, cell density profile, and potential cell migration are all important features to tissue compression stiffening that should be explored further in model tissue systems.

## 5. Conclusions

In this article, we investigated the dynamic meso-scale processes by which a fiber and fiber-bead network remodel under uniaxial compression. We found that compression of fiber networks generates a complex mechanical response that proceeds through a series of remodeling processes, including formation of a densified front, lateral stretch in the network, and fluid flows out of the network. When compression is released, the network relaxes, but with the inclusion of stiff beads, this relaxation is hindered, consistent with a build-up of internal stress and stiffening of the network. Our experiments are thus far the strongest experimental evidence of correlated motion and lateral stretching of a fiber-bead network under compression, consistent with fiber and network stretching predicted to stiffen the network. Most models of fiber network under compression focus on either the formation of a densified front or on fiber network stretching due to lateral stretch or non-affine deformations. These models focus on quasi-equilibrium compression without dynamics. Our experiments highlight the need to consider the effects of poro-elasticity, fiber buckling, and creep in building predictive models that quantitatively capture compression-stiffening in fiber-bead networks, which mimic the compression properties of real tissues. These interactions and their coupled feedback are crucial to many outstanding issues in engineering, physics, and medicine, such as the design of biomedical material to replace damaged tissue. Finally, our work emphasizes the need to study local, dynamic remodeling of networks, where complex material properties dramatically alter the macroscopic mechanical behavior.

## Acknowledgements

We thank Paul Janmey, J.M. Schwarz, and Arvind Gopinath for useful discussions. This work was supported by the *National Institute of General Medical Sciences* of the National Institutes of Health under award number 1R35GM142963-01.

