## Supplementary Information for "Dynamic remodeling of fiber networks with stiff inclusions under compressive loading"

**Supporting Information**

Video S1 Fiber Network (0-20% compressive strain)

Video S2 Fiber-Bead Network (0-20% compressive strain)

Video S3 Fiber Network (20-40% compressive strain)

Video S4 Fiber-Bead Network (20-40% compressive strain)

Video S5 Fiber Network Edge (0-20% compressive strain)

Video S6 Fiber-Bead Network Edge (0-20% compressive strain)

**Video S1-4: Compression Videos.** All videos are 15 frames per second. The compression protocol is compression ramp for ~90s, followed by ~200s of no compression (to allow network to relax). Videos S1-S4 are imaged near the center of the hydrogel. Videos S5-S6 are imaged at the edge of the hydrogel.

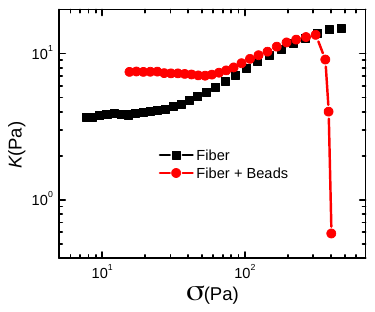

**Figure S1: Differential Shear Modulus vs Shear Stress:** Plot showing the differential shear modulus $K={d\sigma}/{d\gamma}$ vs shear stress $\sigma$ for the fiber and fiber + beads cases. The differential modulus is a better measure for the shear strain stiffening that occurs in the nonlinear regime. The increase in $K$ for the fiber and fiber + beads case corresponds to shear strain stiffening. These values plotted are from the shear strain amplitude sweep data in Figure 1D.

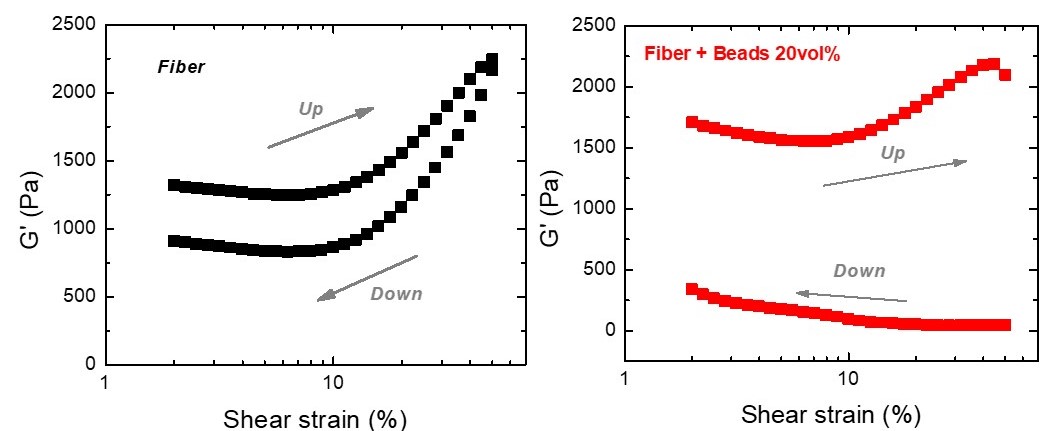

**Figure S2: Shear strain amplitude hysteresis test** Plot showing shear strain amplitude hysteresis, with increasing strain magnitude (up) followed by decreasing strain amplitude (down). In both fiber and fiber + beads cases, there is noticeable hysteresis. With increasing shear strain amplitude, the fiber + beads case has a downturn in stiffness at high shear strain (~40%), similar to that shown in Figure 1d. The descending amplitude sweep is greatly reduced in stiffness for the fiber + beads case. The fiber case, which has no downturn at high shear strain, still exhibits hysteresis.

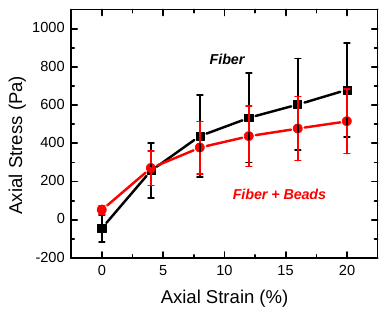

**Figure S3:** **Axial stress vs axial strain for fiber and fiber-bead networks** Plot showing the axial stress vs axial strain values corresponding to the shear modulus data shown in Figure 1e. These values can be used to calculate the Young’s Modulus *E*. The apparent Poisson’s ratio calculated from $E=2G(1+\nu)$ from the above data gives the following values: ν = 3.8 for the fiber case and ν = 1.3 for the fiber + beads case. These unusual values for the Poisson’s ratio indicate the fibrin and fibrin-bead networks are not isotropic linearly elastic materials. Given the large error bars in axial stress, the meaning of the calculated Poisson’s ratios is inconclusive.

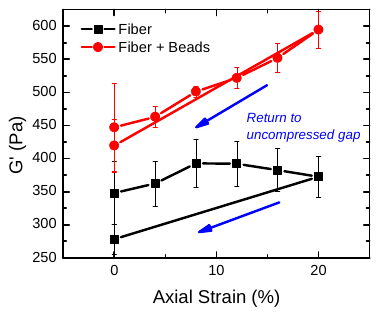

**Figure S4: Compression test hysteresis** Plot of the compression data from Figure 1e with an additional point after the compression test when gap height is returned to the uncompressed state (indicated with blue arrows). Returning to the uncompressed state, the fiber network (black) has a reduction in stiffness from its initial value (~350Pa to ~275Pa), demonstrating a compression test hysteresis. The fiber + beads case (red) only has a minor reduction in stiffness from its original uncompressed value.

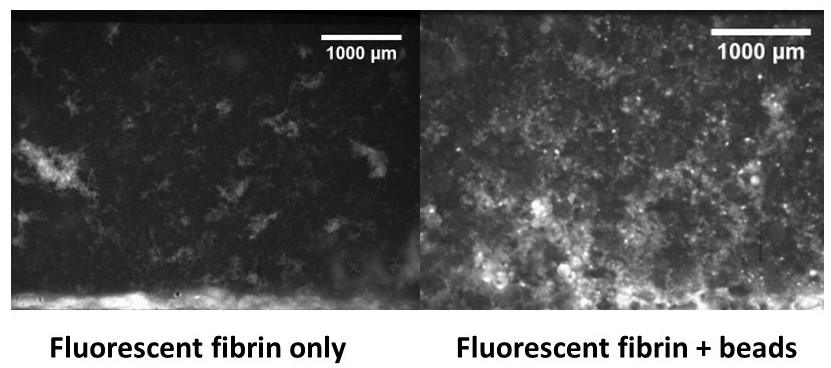

**Figure S5:** **Fluorescently labelled fibrin networks also reveal compression fronts.** Representative images of fluorescently labelled fibrin networks at 40% compressive strain. The compression front formed is similar to the behavior seen when using independent 5micron fluorescent particles in the network. As in the case with fluorescent tracers, the fibrin-bead network reveals a larger front than the fibrin only case.

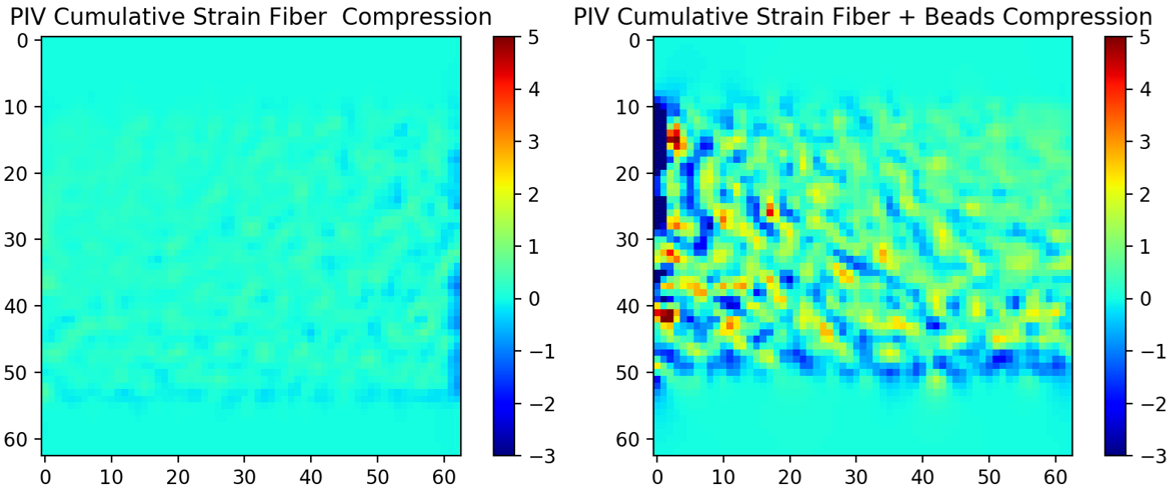

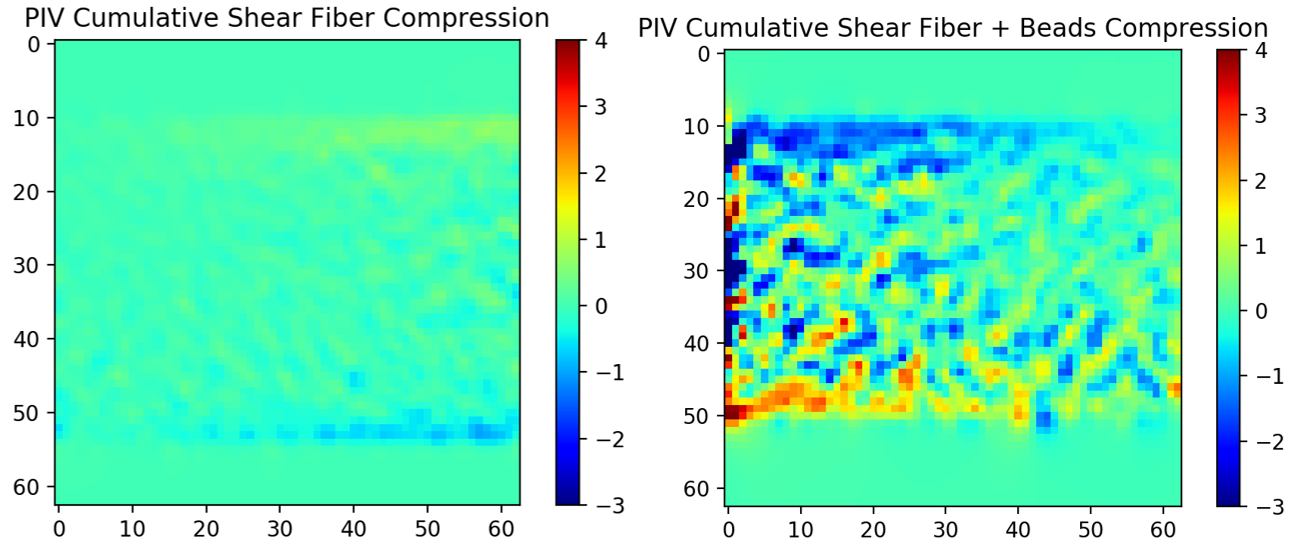

**Figure S6: Cumulative strain and shear maps during compression** Heatmaps demonstrating the cumulative strain (top) and shear (bottom) for the fiber and fiber + beads cases during compression. These images are the accumulated strain and shear during compression up to t = 70s, corresponding to the time of the velocity magnitude maps in Figure 4b.The x, y axis numbering corresponds to the nodal position in Figure 4b.

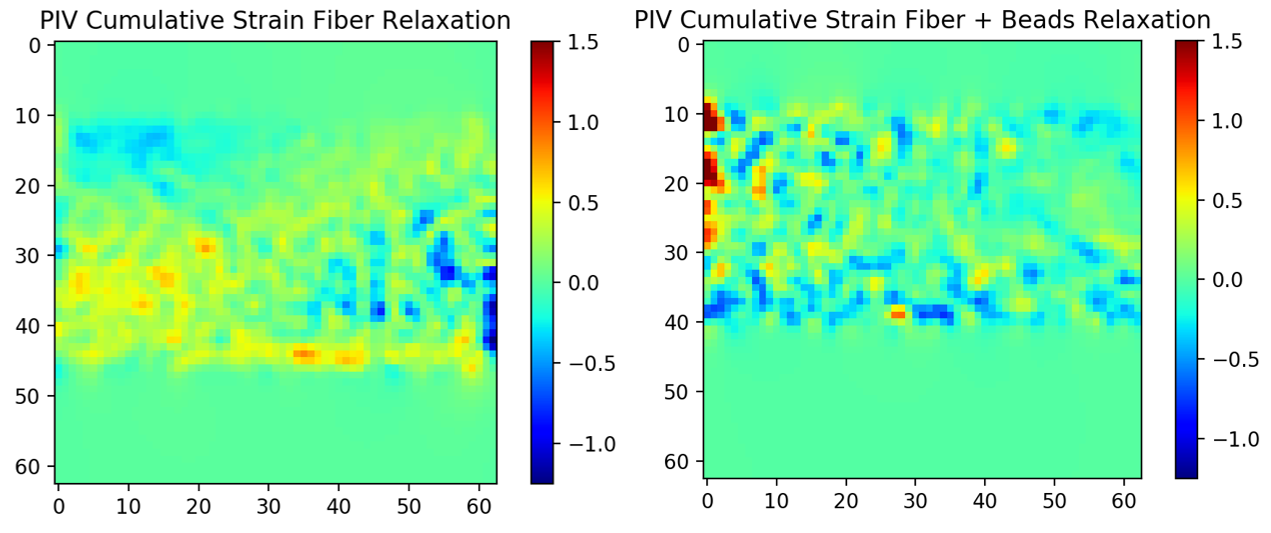

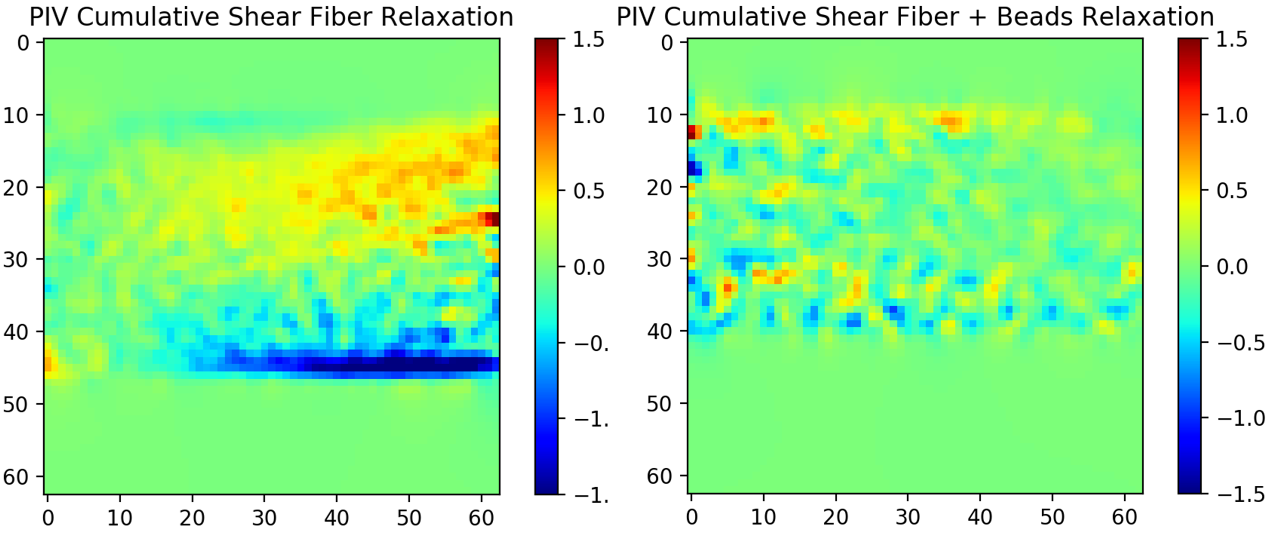

**Figure S7: Cumulative strain and shear maps during relaxation:** Heatmaps demonstrating the cumulative strain (top) and shear (bottom) for the fiber and fiber + beads cases during relaxation. These images are the accumulated shear and strain during the first 20s of relaxation, or t = 110s after the start of the compression, corresponding to the time of the velocity magnitude maps in Figure 4c.The x, y axis numbering corresponds to the nodal position in Figure 4c.

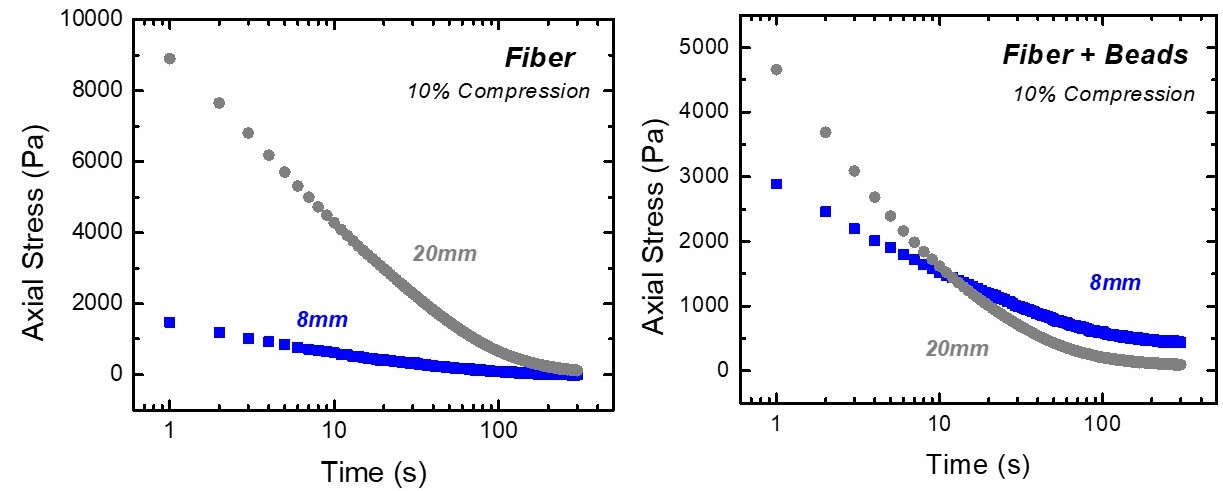

**Figure S8: Axial stress relaxation for two different contact diameters** Plots of the axial stress relaxation at 10% compression for two different rheology geometries: 8mm and 20mm (the geometry used throughout the manuscript). The axial stress relaxation contains contributions from both viscoelasticity and poroelasticity, where the poroelastic contribution is contact area dependent and the viscoelastic is not. For both the fiber and fiber-bead network, there is clearly a distinct axial stress relaxation response for different geometries, indicating the importance of poroelasticity to the relaxation response of this system.
